# Anoxia selects for high fitness biofilms and increases antibiotic resistance in Pseudomonas aeruginosa

**DOI:** 10.64898/2026.07.09.737528

**Authors:** S Bridwell, M. Bahu, C. Okuagu, C.W. Marshall

## Abstract

Antibiotic resistance is a growing global health crisis, yet resistance is almost exclusively quantified under aerobic laboratory conditions that fail to reflect the complex microenvironments bacteria encounter during infection. Many clinically important infection sites, such as airways of individuals with cystic fibrosis or chronic wounds, are microaerobic to anoxic. To address this, we investigated how anoxia alters antibiotic resistance determinants, hypothesizing that anaerobic metabolism might change the fitness effects and selection of resistance mutations. We used experimental evolution to propagate *Pseudomonas aeruginosa* populations for approximately 200 generations under conditions differing in oxygen availability (oxic vs. anoxic), growth mode (biofilm vs. planktonic), and tobramycin (TOB) exposure (subinhibitory increasing to inhibitory concentrations). Subinhibitory exposure was sufficient to achieve resistance 2–4× greater than ancestral levels, with anoxic populations consistently having higher minimum inhibitory concentrations than oxic populations. While resistance mutations in *fusA1* and *ptsP* arose across all conditions, mutations in *amgS* were only selected in oxic populations – indicating condition-specific resistance mutations. Notably, *mexT* mutations were observed in anoxic or tobramycin-exposed populations. The presumed inactivation of *mexT* may also enhance virulence through altered quorum sensing and increased rhamnolipid production. Anoxic populations additionally exhibited significantly increased biofilm formation, reduced twitching motility driven by type IV pilus gene mutations, and greater competitive fitness. Together, these findings demonstrate that oxygen availability shapes resistance evolution in *P. aeruginosa*, with the anoxic environment selecting for a more virulent, sessile, and antibiotic-resistant phenotype.

**Importance:** *Pseudomonas aeruginosa* is an opportunistic pathogen capable of acute and chronic infections that develop antibiotic resistance. One aspect of *P. aeruginosa* that makes it so challenging to treat is its metabolic versatility. We wanted to understand how growth in different infection-relevant conditions, specifically anoxia and surface attachment, would alter the evolutionary pathways of antibiotic resistance. We found that the mutations causing resistance to the commonly used antibiotic tobramycin were condition dependent. We also observed that adaptation to anoxia resulted in *P. aeruginosa* populations that had high biofilm forming capacity and were highly fit compared to its ancestor. This indicates that anoxic infection environments can lead to *P. aeruginosa* variants with increased resistance, increased recalcitrance, and potentially more virulence.

## Introduction

The evolution of antibiotic resistance is an increasingly concerning global problem that complicates the treatment of infections and alters routine medical procedures. In 2019 alone, it is estimated that approximately 1.27 million deaths worldwide were directly related to antimicrobial resistant bacterial infections (“Global Burden of Bacterial Antimicrobial Resistance in 2019,” 2022). A 67.5% increase in antimicrobial resistance related deaths is projected by 2050, rising to 1.91 million per year (Naghavi et al., 2024). The prevalence and recalcitrance of an infecting pathogen is in part due to the fitness effects of the resistance determinants. Bacterial fitness, defined here as the relative ability to survive and reproduce in a specific environment, is a critical metric when assessing the treatment and potential spread of an antibiotic resistant bacterium (Andersson & Hughes, 2010; Melnyk et al., 2015). Furthermore, it has been proposed that the fitness effects of resistant bacteria should be a central factor in assessing the efficacies of potential new drug candidates or treatments (Sommer et al., 2017). Unfortunately, *in vitro* measurements of fitness can be challenging and often don’t accurately reflect either the competitive or the host environment.

Experimental evolution (sometimes referred to as adaptive laboratory evolution (LaCroix et al., 2017)) is a method of studying adaptation in populations exposed to a relatively controlled set of selective pressures (Kawecki et al., 2012; Lenski et al., 1991). This approach allows us to understand resistance mutations that are more fit than the high number of possible resistance mutations in the genome (Schurek et al., 2008) due to competition among coexisting mutations in the population (Cooper, 2018). As a resistance mutation rises to high frequency in a population over time, it is almost certainly adaptive in that selective environment (Cooper, 2018; Lieberman et al., 2011). Therefore, experimental evolution can be a powerful approach to understand low-cost and potentially high fitness antibiotic resistance mutations in a set of defined environmental conditions and selective pressures (Ahmed et al., 2018; Santos-Lopez et al., 2019).

This approach can be particularly useful when trying to tease apart the relative contributions of selective pressures from complex environments like the human infection environment. When an infection is established, the nutrients are often changing (Hendricks et al., 2016; La Rosa et al., 2019), the immune system deploys defenses (Moser et al., 2021), and oxygen depletion can rapidly occur (Cowley et al., 2015; Worlitzsch et al., 2002). We are particularly interested in the latter, where oxygen depletion can alter antibiotic resistance outcomes. It has been demonstrated that many infection environments are microaerophilic or anaerobic (Cowley et al., 2015; Hassett et al., 2009), and that anaerobic conditions increase the resistance levels in bacteria to antibiotics (Borriello et al., 2004; Hill et al., 2005; Walters et al., 2003). We therefore tested how the intersection between antibiotic exposure and oxygen depletion altered the evolutionary pathways of resistance in the facultatively anaerobic opportunistic pathogen *Pseudomonas aeruginosa*.

*Pseudomonas aeruginosa* is a Gram-negative, ubiquitous pathogen with the ability to grow in both aerobic and anaerobic conditions, using oxygen or nitrate as a terminal electron acceptor for respiration (Hassett et al., 2009). It causes nosocomial infections across healthcare settings (Rosenthal et al., 2016) as well as chronic infections including pneumonia in people with cystic fibrosis (CF) (Lyczak et al., 2002). *Pseudomonas aeruginosa* is considered a serious threat level pathogen by the CDC (Centers for Disease Control and Prevention (U.S.), 2019), and a high priority pathogen by the WHO (*WHO Bacterial Priority Pathogens List 2024*, 2024), due to the rapid antibiotic resistance development and established multidrug resistance. One mechanism of oxygen depletion, and perhaps one of the earliest adaptations by *P. aeruginosa* in infection (Gloag et al., 2019), is the prolific biofilm-forming capabilities of *P. aeruginosa* (Høiby et al., 2010; Soares et al., 2020). Biofilms are groups of surface-attached cells encased in a matrix, which can be made up of extracellular DNA, proteins, and polysaccharides (Flemming & Wingender, 2010), and in some cases filamentous phage (Secor et al., 2015). Because of this protective matrix, slow diffusion, and the reduction of metabolic activity in biofilms, biofilm communities are often difficult to eradicate (Høiby et al., 2010; Stewart & Costerton, 2001). Therefore, we wanted to understand how biofilms and the absence of oxygen could alter antibiotic resistance pathways in *P. aeruginosa*.

To test this intersection of biofilm and oxygen depletion on antibiotic resistance pathways, we used the antibiotic tobramycin (TOB), an aminoglycoside antibiotic commonly used to treat infections and a key anti-*P. aeruginosa* treatment in cystic fibrosis (CF) (Mogayzel et al., 2013). Concentration gradients can be created in the CF airways with TOB usage, as the antibiotic does not always penetrate the thick mucus layer and biofilms in the airway (I. Martin et al., 2021; Rouillard et al., 2024). Bacteria residing deeper in the mucus or biofilm may experience subinhibitory concentrations of antibiotic, which has the potential to drive mutational resistance (Gullberg et al., 2011; Ramsay et al., 2021; van der Horst et al., 2011; Wistrand-Yuen et al., 2018). Many niches of the CF airway are anaerobic (Cowley et al., 2015; Worlitzsch et al., 2002), and the combination of subinhibitory antibiotic exposure and reduced oxygen represents a clinically relevant selective setting that has not been investigated through experimental evolution.

The goal of the current study was to determine how different selective pressures common in the infection environment interact to confer resistance and to quantify the resulting fitness effects. We hypothesized that anaerobic environments would lead to antibiotic resistance mutations distinct from those selected under aerobic conditions. The hypothesis was tested by using experimental evolution where we propagated *P. aeruginosa* for approximately 200 generations in both the presence and absence of oxygen, in either a biofilm or planktonic growth mode, and with and without tobramycin (TOB) exposure. We also tested how the resulting adaptation to biofilm growth and anaerobic conditions altered phenotypes and the fitness of evolved populations relative to the ancestor, offering insight into the adaptive value of the acquired mutations in each selective condition.

## Results

To understand how different lifestyles could alter antibiotic resistance evolution, *P. aeruginosa* strain MPAO1 was used to inoculate six replicate lineages for each of eight conditions including oxic or anoxic growth, biofilm or planktonic growth, and with or without the antibiotic tobramycin (TOB) (**Fig 1**) (Poltak & Cooper, 2011; Santos-Lopez et al., 2019; Traverse et al., 2013). Populations were propagated for 30 days using a two-step experiment where TOB was increased: 15 days at 0.5X the minimum inhibitory concentration (MIC) then another 15 days at 1X ancestral MIC.

**Fig 1.**
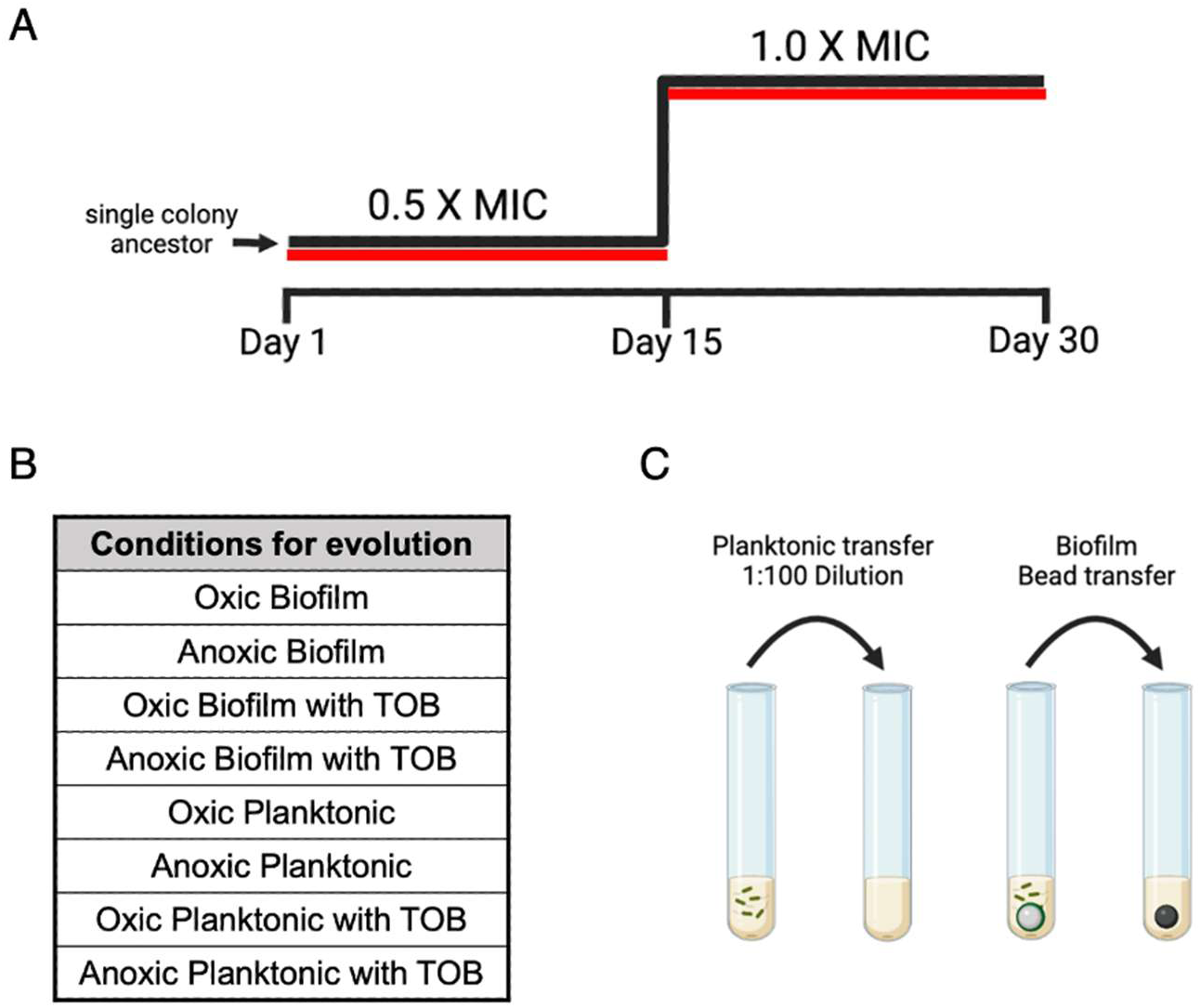
Populations of *P. aeruginosa* were propagated in minimal M9 media for 30 days. (A) Populations were exposed to a subinhibitory (0.5X MIC) concentration of tobramycin for the first 15 days, with the concentration being doubled (1X MIC) for the second half of the experiment. Six replicate populations per condition were propagated. Populations were saved throughout the experiment for sequencing and phenotypic testing. (B) Populations were evolved under eight conditions: alternating oxic or anoxic, biofilm or planktonic, and with tobramycin or without tobramycin. (C) Populations were propagated with either planktonic growth in a daily 1:100 dilution or biofilm growth with a daily biofilm bead transfer (Poltak & Cooper, 2011).

### Subinhibitory tobramycin exposure decreased susceptibility

All populations exposed to subinhibitory concentrations of TOB achieved elevated resistance relative to the MPAO1 ancestor (**Fig 2**). Ancestral MIC values ranged from 1-2 mg/L when measured in oxygen and 2-4 mg/L when measured under anoxic conditions. Populations propagated without TOB exposure showed no substantial increase in resistance at either timepoint, demonstrating that antibiotic exposure, rather than the environmental conditions alone, drove the MIC increases.

**Fig 2.**
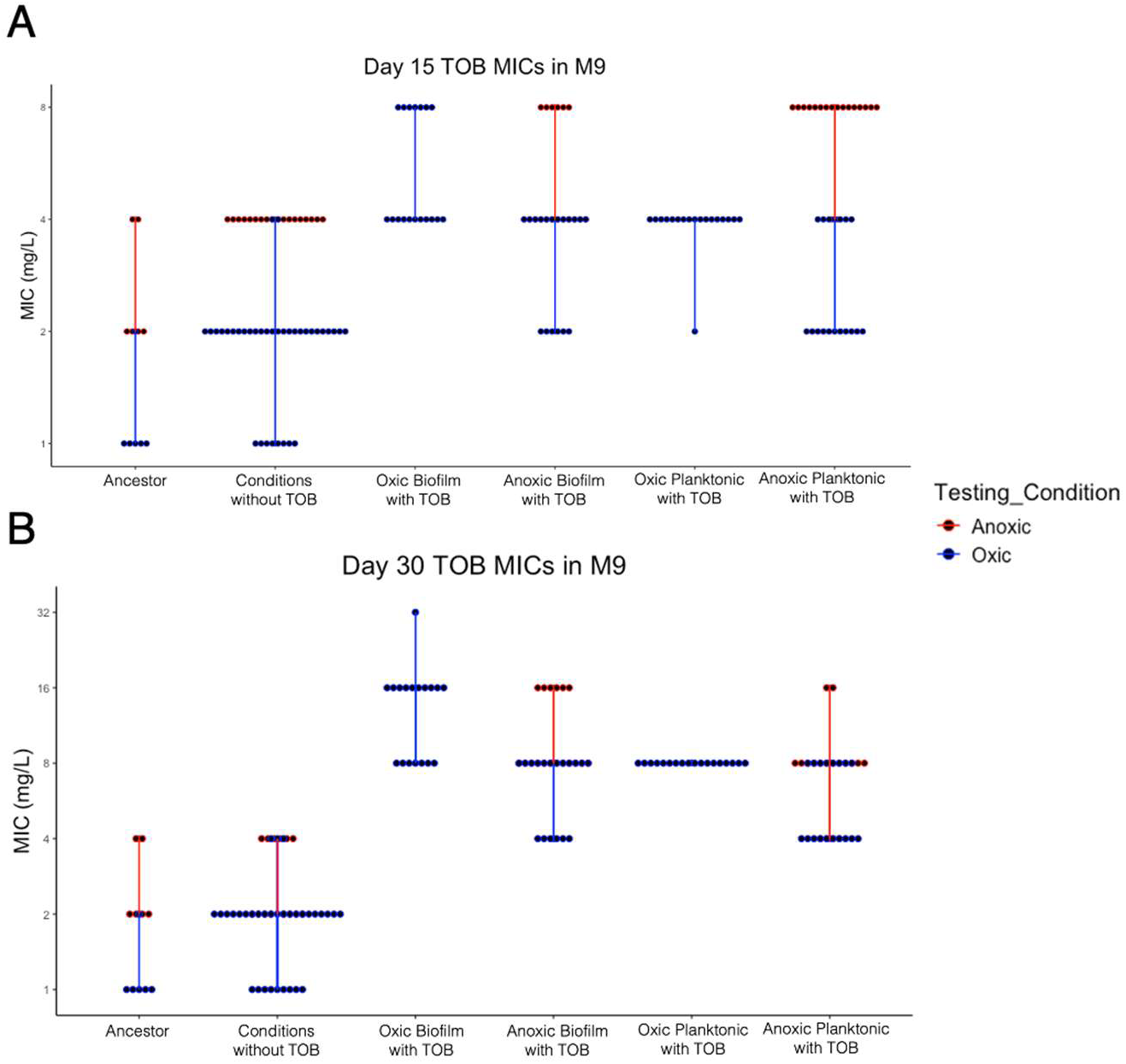
Minimum inhibitory concentrations (MICs) of evolved populations following 15 days of subinhibitory exposure of tobramycin (A) and after 15 additional days at the inhibitory concentration (B). MICs were done by broth microdilution in minimal M9 media under oxic growth (in blue) or anoxic growth (in red). Three technical replicates per population are shown. MICs for populations propagated under conditions lacking TOB exposure were grouped. The ancestral clone MIC was between 1-2 mg/L in minimal M9 media. Populations needed to gain resistance to TOB to survive the second half of the experiment with TOB exposure at the concentration of the MIC (1X MIC).

After 15 days of subinhibitory TOB exposure, all TOB-exposed populations had already achieved elevated MICs relative to the ancestor (**Fig 2A**). This shows that clinically relevant resistance emerged rapidly under subinhibitory antibiotic pressure alone. By day 30, following an additional 15 days at 1X MIC, resistance levels increased further across all TOB-exposed conditions (**Fig 2B**). Both oxic and anoxic TOB-exposed populations achieved MICs of at least 2-4-fold above the ancestral baseline, with several populations reaching the MIC values at or above the clinical susceptibility breakpoint for TOB (≥8 mg/L; (Clinical and Laboratory Standards Institute, 2026). However, the breakpoint levels of resistance were measured in the experimental minimal media M9+ instead of the designated standard media Mueller-Hinton broth, where MICs were higher than the ancestor but not quite at the clinical breakpoint (**Fig S1**). Notably, MICs measured anoxically consistently displayed higher MICs compared to when measured in the presence of oxygen (**Fig 2**). Together, these data show that even low-level antibiotic exposure is enough to decrease susceptibility, potentially to a level of clinical significance and that although not often considered in susceptibility testing, oxygen availability in an environment can change the degree of resistance.

### Condition-dependent genomic targets of TOB resistance

To identify mutations associated with lifestyle and TOB resistance phenotypes, whole population sequencing was performed on evolved populations on day 30 (after exposure to 1X the ancestral MIC). Four evolutionary replicates per TOB-exposed condition and two evolutionary replicates per no TOB-exposed condition were sequenced and compared with the ancestor. TOB resistance across conditions was predominantly associated with mutations in *amgS*, *fusA1*, and/or *ptsP* (**Fig 3, Table S1)**. Each of these genes has previously been implicated in aminoglycoside resistance in *P. aeruginosa* in both clinical and laboratory studies (Bolard et al., 2018; Lau et al., 2013; Scribner et al., 2020). *fusA1* encodes elongation factor G (EF-G), an essential GTPase that facilitates translocation during protein synthesis (Rodnina, 2013). Mutations in EF-G have been previously indicated in TOB resistance (Bolard et al., 2018; Scribner et al., 2020), and have been repeatedly identified in longitudinal clinical isolates as a convergently evolved resistance locus (López-Causapé et al., 2017; Marvig et al., 2015). *ptsP* encodes the phosphoenolpyruvate-protein phosphotransferase enzyme of the nitrogen phosphotransferase system (PTS) (Pflüger-Grau & Görke, 2010) and is known to increase TOB resistance in *P. aeruginosa* (Abisado et al., 2021; Scribner et al., 2020). Selection for *fusA1* and *ptsP* mutations across all TOB-exposed conditions suggests that these resistance mutations confer a relatively high fitness solution to resistance across the varied environments.

**Fig 3.**
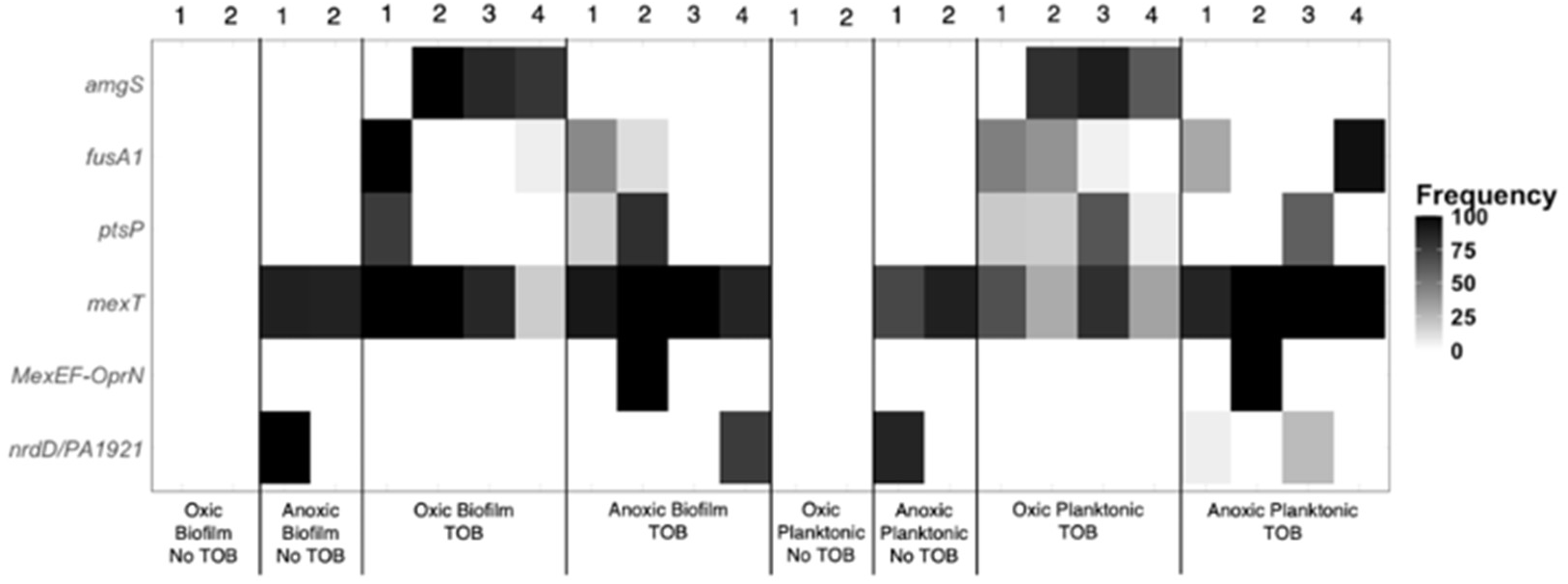
Mutation identification and frequency determined by whole-population genome sequencing of each evolution condition of *P. aeruginosa*. Four replicates per TOB-exposed condition and two replicates per condition without TOB exposure from day 30 of the evolution were selected for sequencing. Shading intensity indicates total frequency of mutations in noted genes within a population.

Mutations in *amgS* were notably enriched in oxic-evolved, TOB-exposed populations, consistent with a condition-dependent resistance mechanism. The *amgS* gene encodes the sensor histidine kinase of the AmgRS two-component system, which is involved in combating aminoglycoside-induced membrane stress by upregulating genes involved in membrane protein quality control (Lau et al., 2013; S. Lee et al., 2009). This pattern of condition-dependent antibiotic resistance shows that the mechanism of resistance is not fixed but can be shaped by antibiotic and environmental pressures.

### Selection for mutations in *mexT*

Interestingly, mutations in *mexT*, a LysR-type transcriptional activator of the *mexEF-oprN* efflux pump operon, were found at high frequency in all conditions either exposed to TOB or exposed to anoxia but not in the oxic conditions without TOB (**Fig 3**). We observed mostly point mutations throughout the *mexT* gene (**Fig S2, Table S2**), but we also observed a few large deletions and two populations had deletions of the entire *MexEF-OprN* operon, suggesting strong selection against expression of this efflux system. The occurrence of *mexT* mutations and independent large deletions across replicates and conditions is a strong indicator of selection for gene inactivation. MexT increases expression of MexEF-OprN, which is an RND-family pump used in export of fluoroquinolones, chloramphenicol, and trimethoprim antibiotics, but not known to efflux aminoglycosides (Köhler et al., 1997; Llanes et al., 2011). We tested chloramphenicol susceptibilities in populations with *mexT* mutations and found that populations with these mutations had increased susceptibility to chloramphenicol, indicating loss of function mutations that inactivate or repress expression of the MexEF-OprN efflux pump (**Fig S3**). Despite evidence that suggests the MexEF-OprN efflux pump does not use TOB as a substrate (Köhler et al., 1997; Llanes et al., 2011; Masuda et al., 2000), mutations in *mexT* have been shown to decrease aminoglycoside susceptibility, possibly through alterations in the LPS layer of the cell surface (Figueroa et al., 2025). Recently, *mexT* deletion mutants were shown to downregulate O-antigen biosynthesis resulting in a modification in the LPS layer of *P. aeruginosa* that would interfere with aminoglycoside entry into the cell (Figueroa et al., 2025). Because *mexT* mutations were also found in anoxic populations with no TOB exposure, their presence likely indicates a fitness benefit outside of the antibiotic exposure. MexT is a transcriptional regulator whose activation influences multidrug resistance, virulence, metabolic, and quorum sensing functions (Kostylev et al., 2023; Tian, Fargier, et al., 2009; Tian, Mac Aogáin, et al., 2009). Because of its role in many of these functions, *mexT* loss of function mutations have been identified in clinical *P. aeruginosa* isolates from CF samples (Sherrard et al., 2017; Smith et al., 2006). In our MPAO1 strain, *mexT* is thought to be constitutively active and has been observed during serial passage to revert to a phenotype that inactivates *mexT*, often through loss of function mutations (S. Lee et al., 2021; Luong et al., 2014). It is curious that these putative loss of function *mexT* mutations only occurred in our study when populations were propagated either in the presence of TOB or when propagated anaerobically (**Fig 3**). No *mexT* mutations were observed in any of the aerobic no-TOB conditions, indicating in our study that these mutations were not generally adaptive but only adaptive under anaerobic or antibiotic conditions.

### Anoxic adaptations drive enhanced biofilm formation

Biofilm formation is a significant feature of chronic *P. aeruginosa* infection in the CF lung, promoting the persistence of infections even under host immune response and antibiotic treatment (Bjarnsholt et al., 2009; Høiby et al., 2010). We therefore wanted to understand how adaptation to different lifestyles would affect biofilm formation.

Populations evolved under anoxic conditions, particularly the anoxic biofilm populations propagated without TOB, showed an increase in biofilm formation when tested under anoxic conditions (**Fig 4**). Biomass measured over 1000% of anoxic ancestor control levels in the anoxic, no TOB populations (p <0.001, Dunnett’s test). Populations evolved under oxic conditions, on the other hand, showed relatively modest changes in biofilm formation under both oxic and anoxic testing conditions. These comparisons demonstrate that the anoxic selective pressure and not prolonged laboratory propagation as a biofilm, was the primary driver of the increased biofilm levels. The significantly enhanced anoxic biofilm phenotype indicates that adaptation to an anoxic environment selects for physiological or regulatory changes that promote biofilm formation under no oxygen conditions, potentially showing specialization directly applicable to chronic infections (Høiby et al., 2010; Worlitzsch et al., 2002).

**Fig 4.**
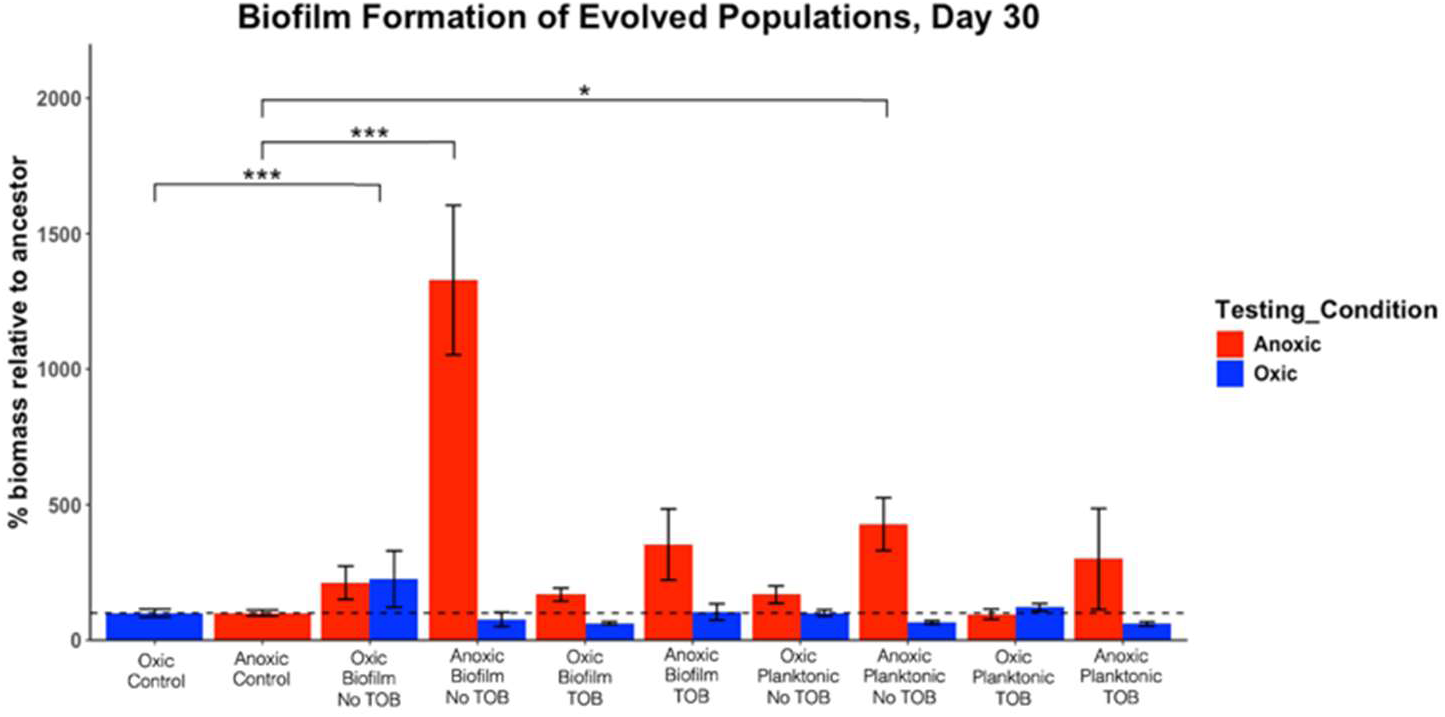
Biofilm formation for all conditions was grown under both oxic (blue) and anoxic (red) testing conditions. Results are shown relative to the ancestor with ancestor values set at 100%. Oxic measurements were compared with the oxic control and anoxic measurements were compared with the anoxic control. A one-way ANOVA (p<0.05) and Dunnett’s post hoc test were done to compare anoxic testing condition values to the anoxic control and oxic testing condition values to the oxic control (Dunnett’s test: *= p<0.05, ***= p<0.001).

### Mutations in type IV pilus genes are enriched and alter twitching motility

Type IV pili (T4P) are retractile surface appendages that are critical in the motility, biofilm formation, pathogenesis, and host colonization of *P. aeruginosa* (Burrows, 2012). T4P expression and twitching motility are gradually lost over time during chronic infection, transitioning from acute, motile phenotypes to a sessile, biofilm lifestyle (Cullen & McClean, 2015; Smith et al., 2006). Because our anoxic-evolved populations developed enhanced biofilm phenotypes, we predicted that motility would also be altered. Multiple evolved populations showed significantly different motility relative to the ancestor (**Fig 5A**). Reductions in motility below the ancestral level were observed across multiple conditions but were most prevalent and consistent in anoxic planktonic and to some extent anoxic biofilm populations evolved with TOB. Oxic evolved populations were generally at or above ancestor motility levels, though some reductions were observed in oxic populations as well. The decrease in twitching motility observed in those anoxic, TOB-evolved populations is consistent with the phenotypic switch from type IV pili-based motility to increased biofilm formation observed in many chronic infections (Burrows, 2012; B. Lee et al., 2005).

**Fig 5.**
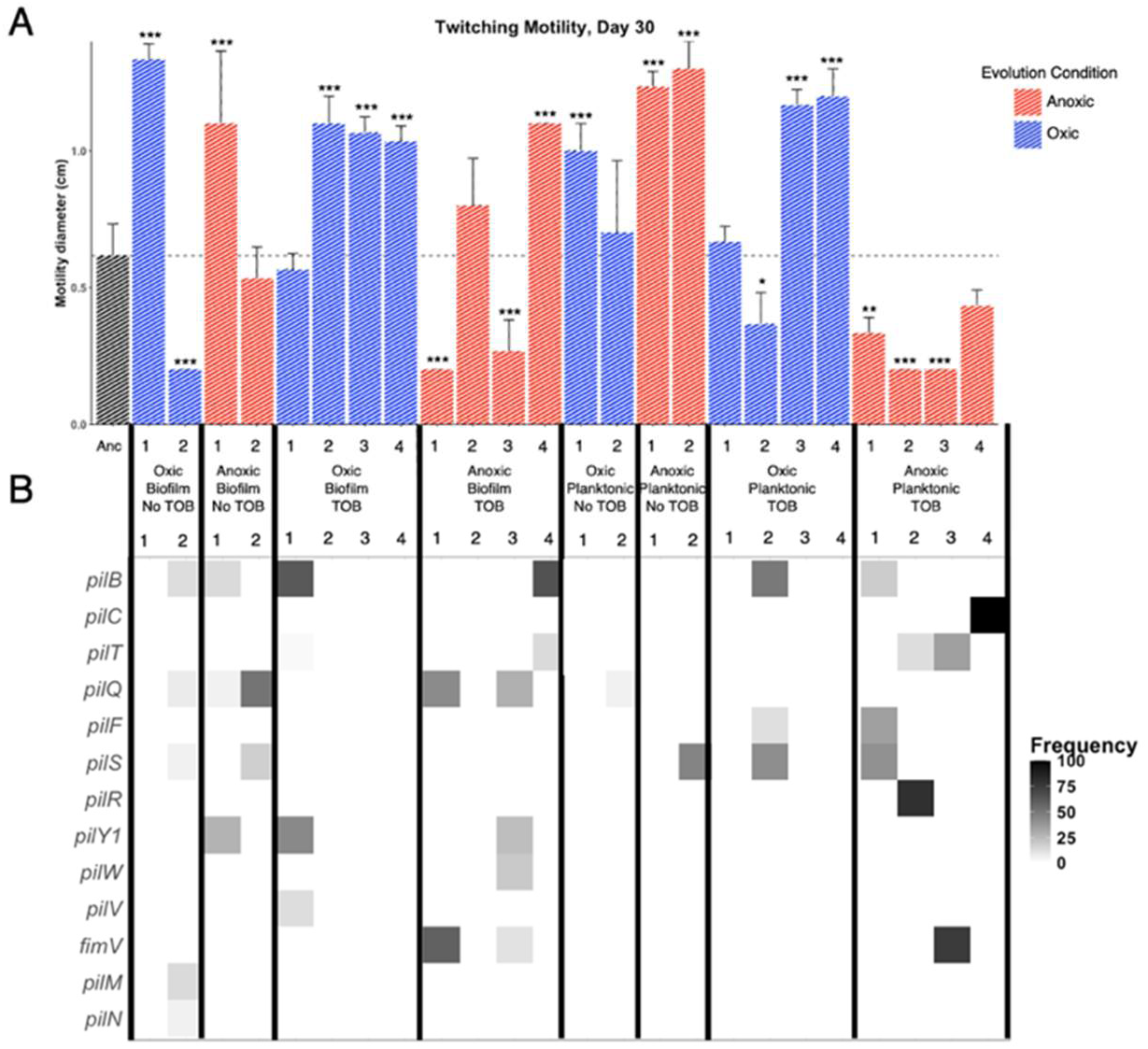
Twitching motility (A) and mutations (B) associated with type IV pilus genes. A one-way ANOVA (p<0.05) and Dunnett’s post hoc test was performed to compare ancestor motility to motility of evolved populations (Dunnett’s test: *=p<0.05, **=p<0.01, ***=p<0.001). (B) Mutation identification and frequency determined by whole-population genome sequencing of each evolution condition of *P. aeruginosa*. Four replicates per TOB-exposed condition and two replicates per condition without TOB exposure from day 30 of the evolution were selected for sequencing. Shading intensity indicates total frequency of mutations in noted genes within a population.

Accompanying the phenotypic changes in twitching motility, mutations in genes encoding T4P assembly, regulation, and retraction machinery were observed in all conditions (**Fig 5B**). These included structural components (*pilB, pilC, pilM, pilN*), assembly-associated genes (*pilF, pilW, pilV*), the two-component regulatory system (*pilS, pilR*), the anti-retraction factor *pilY1*, the retraction ATPase *pilT*, and the related scaffold protein *fimV* (Leighton et al., 2015). Mutations in these genes were more prevalent in anoxic populations (22 mutations compared to 12 in oxic populations) and there were significantly more T4P mutations in populations with lower twitching motility diameter compared to populations with higher or no change in motility (Mann Whitney U, p<0.05), however the phenotypic effects of these mutations were nuanced as certain genes and combinations of mutations can either increase or decrease twitching motility (Burrows, 2012; Leighton et al., 2015). We did test motility in transposon-insertion mutants disrupting either *fimV*, *pilB*, *pilQ*, or *pilT* and found that each of these mutants was deficient in twitching motility (**Fig S4**). Taken together, these findings support the relevance of our model and provide insight into selective conditions favoring T4P mutations similar to *in vivo* studies where mutations in T4P genes are often found (Cullen & McClean, 2015; Marvig et al., 2015; Smith et al., 2006).

### High competitive fitness of anoxic evolved lineages

Bacterial fitness is an important metric for understanding the clinical relevance of antibiotic resistance (Melnyk et al., 2015; Sommer et al., 2017). Across all anoxic-evolved populations tested, fitness tested under anoxic conditions was significantly higher relative to the anoxic control (p<0.01-0.001, Dunnett’s test), while fitness under oxic conditions was largely unchanged (**Fig 6**). The oxic-evolved populations did not have reciprocal oxic fitness gains, demonstrating that adaptation to anoxia alone rewires *P. aeruginosa* for high competitive fitness. Furthermore, at high selection rates where r>2, it is possible that the anoxic-evolved populations are actively inhibiting the ancestor rather than outcompeting it for resources. Assigning genotypic explanations for the high fitness phenotypes can be challenging because of the difficult to predict epistatic interactions between mutations, but presumed loss of function mutations in *mexT* were one commonality between all anoxic-evolved populations (**Fig 3**) that might lead to increased anoxic fitness. In support of this, we observed increased fitness in transposon-insertion mutants in *mexT* (**Fig S5**).

**Fig 6.**
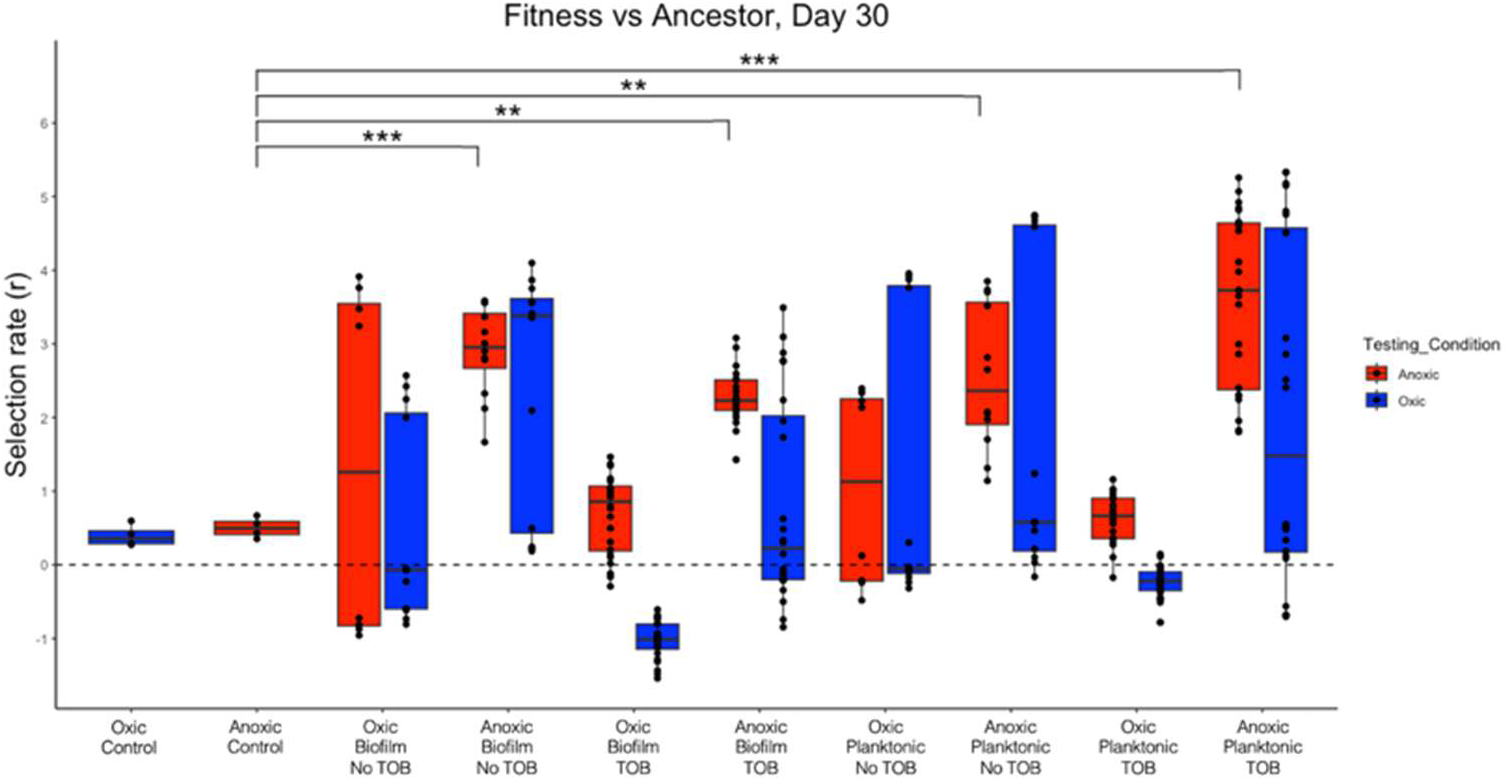
Fitness was evaluated by growing and competing the evolved populations (on x-axis) against the ancestor under both oxic (blue) and anoxic (red) conditions. Fitness was quantified as selection rate (r), value greater than 0 indicates higher fitness in the evolved strain and a value less than 0 indicates higher fitness in the ancestor. A one-way ANOVA (p <0.05) and Dunnett’s post hoc test was performed to compare selection rate of controls to evolved populations (Dunnett’s test:**=p<0.01, ***=p<0.001). Assays tested under oxic conditions were compared with the oxic control and anoxic assays were compared with anoxic control.

Populations evolved under TOB exposure showed a trend toward decreased fitness relative to other evolution conditions, particularly the oxic biofilm condition with TOB exposure. This could potentially reflect the metabolic burden associated with carrying resistance mutations like *fusA1* or *amgS.* The amgS mutations selected in oxic TOB-exposed populations activate the AmgRS membrane stress response, which involves upregulation of membrane protein quality control functions (S. Lee et al., 2009).

## Discussion

Our results highlight several characteristics that may be of clinical significance: (i) subinhibitory TOB exposure is sufficient to increase resistance across all conditions, with anoxic measurements consistently reaching higher MICs; (ii) resistance is achieved through a combination of known resistance alleles, condition-dependent genomic targets, and underreported alleles that alter susceptibility specifically under anoxia; (iii) anoxic evolution selects for high biofilm formation and decreased twitching motility, phenotypes that are characteristic of chronic infection; and (iv) anoxic-evolved populations show increased fitness under anoxic competition, a finding that may have implications in understanding pathogen success in infection environments. These results emphasize the consideration that resistance evolution under conditions similar to the infection environment may uncover adaptive trajectories that would be inaccessible under standard laboratory conditions. *Pseudomonas aeruginosa* is categorized as a serious threat level pathogen (Centers for Disease Control and Prevention (U.S.), 2019) because of its capacity for antibiotic resistance development and prolific ability to form biofilms (Flemming & Wingender, 2010), which can create the oxygen depleted niches that drive the adaptive pathways that we characterize here (Høiby et al., 2010).

Although this context is well-established, most experimental evolution and resistance studies rely on aerobic physiology. The progression of *P. aeruginosa* in cystic fibrosis-related chronic infections is shaped by a thickened mucus layer in the airways that creates oxygen gradients and anoxic niches (Hassett et al., 2009; Worlitzsch et al., 2002). These anoxic niches and presumably prevalent anaerobic metabolism leave a critical gap in our understanding of how oxygen limitation shapes adaptation in an infection.

Consistent with a growing body of literature, we demonstrate that sub-MIC antibiotic exposure selected for resistance through mechanisms distinct from what is identified at lethal antibiotic levels (Gullberg et al., 2011; Pereira et al., 2023; Wistrand-Yuen et al., 2018). Wistrand-Yuen et al. showed that *Salmonella* exposed to sub-MIC streptomycin evolved increased resistance through accumulation of mutations affecting ribosomal RNA, aminoglycoside uptake, and antibiotic modifying enzyme induction. These pathways have not been shown to be selected at inhibitory concentrations, which suggests that weak selective pressure can lead to clinically significant resistance outcomes (Wistrand-Yuen et al., 2018). More recently, Pereira et al. showed that many mutations selected for under subinhibitory concentrations of antibiotic have been found in clinical isolates, reinforcing the relevance of sub-MIC environments that can be a source of clinically important resistance (Pereira et al., 2023). Antibiotic concentration gradients can be generated in the CF airways with TOB usage, as the antibiotic has limited infiltration of the thick mucus layer and biofilms within the airway (Geller et al., 2002; I. Martin et al., 2021) yet is commonly used to treat CF infections (Mogayzel et al., 2013; Ramsey et al., 1999). Bacteria residing deeper in the mucus or biofilm may experience subinhibitory concentrations of antibiotics, which furthers the potential to drive resistance (Ramsay et al., 2021; van der Horst et al., 2011).

Many niches of the CF airway are anaerobic (Cowley et al., 2015; Worlitzsch et al., 2002) and the combination of subinhibitory antibiotic exposure and reduced oxygen represents a problematic setting for resistance evolution that has been understudied. Our finding that 15 days of subinhibitory TOB exposure led to MICs at or above the clinical susceptibility breakpoint (Clinical and Laboratory Standards Institute, 2026) in both oxic and anoxic evolved populations underscores the risk posed by complex infection environments. Additionally, MICs measured under anoxic testing conditions were consistently higher than oxic MICs, regardless of evolutionary condition, suggesting that anoxic physiology increases resistance outcomes and that traditional aerobic susceptibility testing may systematically underrate resistance in infections with anoxic niches (Borriello et al., 2004; Hill et al., 2005; L. W. Martin et al., 2023; Walters et al., 2003).

TOB resistance was not achieved through a single mechanism but through condition-dependent genomic targets. Specifically, the condition-dependent selection of *amgS* mutations in oxic populations compared to their absence in anoxic populations suggests that aminoglycoside resistance is strongly shaped by oxygen availability in an environment. The AmgRS two-component system defends against membrane damage caused by aminoglycosides and has been found in clinical isolates conferring aminoglycoside resistance (Lau et al., 2013; S. Lee et al., 2009). More work is needed to understand why these mutations were advantageous under aerobic conditions but not found when the cells were growing anaerobically.

Across all TOB-exposed populations, reduced susceptibility was seen through mutations predominantly in *fusA1 and ptsP*. *fusA1* and *ptsP* represent well-established routes to aminoglycoside resistance in *P. aeruginosa*. *fusA1* encodes elongation factor (EF-G) which is an essential GTPase of the translational machinery (Rodnina, 2013). Mutations in *fusA1* have been repeatedly identified in CF isolates associated with aminoglycoside resistance (Bolard et al., 2018; López-Causapé et al., 2017). The *ptsP* gene encodes the nitrogen-specific phosphoenolpyruvate phosphotransferase system enzyme I and although not expected to be a direct target of the drug, it has been previously identified as an alternative route for TOB resistance in *P. aeruginosa* (Scribner et al., 2020). The exact mechanism of resistance is not fully understood, but the nitrogen phosphotransferase system has been shown to interact with the GigA/GigB signal transduction pathway that works to direct the global transcriptional response to antibiotic stress in other gram negative pathogens (Gebhardt & Shuman, 2017), with homologous functions implicated in *P. aeruginosa* (Abisado et al., 2021).

The widespread selection for *mexT* mutations across all TOB-exposed and/or anoxic conditions is an important finding that cannot be explained by aminoglycoside efflux. The efflux pump activated by MexT, MexEF-OprN, does not transport aminoglycosides (Llanes et al., 2011), yet *mexT* mutations were found at high frequency in all TOB-exposed populations and in no-TOB anoxic conditions. MexT both activates MexEF-OprN and suppresses virulence through indirect inhibition of the *RhlI-R* quorum sensing system and *Pseudomonas* quinolone signal (PQS) system (Frando et al., 2025; Kostylev et al., 2023). This is likely due to MexEF-OprN mediated export of the PQS precursor 4-hydroxy-2-heptylquinoline (HHQ), which results in reduced production of virulence factors pyocyanin, rhamnolipids, and elastase (Frando et al., 2025; Köhler et al., 2001; Kostylev et al., 2023). Inactivation of *mexT* is therefore expected to reverse these effects, increasing production of these virulence factors and decreasing *mexEF-oprN* expression. Our results are consistent with *mexT* inactivation demonstrated by the highly competitive fitness of the mutant alleles (**Fig 6**)(**Fig S5**) and increased susceptibility to chloramphenicol (**Fig S3**)(Held et al., 2012). The diversity of *mexT* mutations (**Fig S2**)(**Table S1**) across independent replicates is indicative of strong positive selection (Lieberman et al., 2011; Wong et al., 2012). *mexT* loss of function mutations have been documented in clinical CF isolates (Richardot et al., 2016; Sherrard et al., 2017), and our data provide experimental evidence that these arise specifically under anoxic and/or antibiotic selective conditions that mimic the CF airway. This helps to establish *mexT* as a broad adaptive regulator whose inactivation affects antibiotic resistance, virulence, and competitive fitness under anoxic conditions.

Anoxic-evolved populations, particularly anoxic biofilm populations propagated without TOB, exhibited significantly increased biofilm formation when tested under anoxic conditions (**Fig 4**). This enhancement was specific to anoxia as the same populations showed modest or no increase in biofilm under oxic testing and oxic-evolved populations showed little change under either testing condition. This demonstrates that the enhanced biofilm phenotype was a central aspect of the adaptation to anoxia and not a broad adaptation to prolonged laboratory propagation. These results are consistent with the well-recognized link between c-di-GMP signaling, anaerobic gene regulation, and biofilm commitment in *P. aeruginosa* (Römling et al., 2013). Under anoxic conditions, the anaerobic regulator Anr upregulates denitrification, which produces nitric oxide (NO) as an intermediate (Hammond et al., 2015; Toyofuku et al., 2012). At the low concentrations generated endogenously, NO promotes elevated c-di-GMP and biofilm formation rather than dispersal (Cutruzzolà & Frankenberg-Dinkel, 2015; Park et al., 2020). In chronic infections, *P. aeruginosa* populations face antibiotic pressure and oxygen depletion which together favor increased biofilm formation (Borriello et al., 2004; Cowley et al., 2015; Worlitzsch et al., 2002). Increased biomass then further reduces antibiotic penetration and oxygen availability, strengthening the environmental pressures that drove the adaptation in the first place (Borriello et al., 2004; Ciofu & Tolker-Nielsen, 2019; Stewart & Costerton, 2001; Walters et al., 2003).

The T4P mutations enriched in anoxic-evolved populations (**Fig 5B**) reinforce the high biofilm phenotype. Mutations in *pilB*, *pilC*, *pilM*, *pilN*, *pilS*, *pilR*, *pilT*, *pilY1*, and/or *fimV* reflect independent selection for T4P inactivation, leading to a sessile phenotype. *pilT* mutations are particularly notable as PilT is the primary retraction motor and *pilT* mutants can be hyperpiliated, rendering them unable to twitch; this phenotype is known to be associated with biofilm formation (Burrows, 2012). Similarly, *fimV* encodes a peptidoglycan-binding protein needed for T4P assembly. *fimV* mutations disrupt biogenesis of the pilus through impaired PilQ secretin formation and assembly (Wehbi et al., 2011). The gradual loss of twitching motility is also documented in longitudinal CF isolates as infections progress (Cullen & McClean, 2015; Marvig et al., 2015; Smith et al., 2006), possibly connecting that clinical trajectory directly to anoxic and biofilm selective pressures.

Anoxic-evolved populations consistently outcompeted the ancestor under anoxic conditions across all evolution conditions tested, with selection rates significantly above zero and often above 2 (**Fig 6**). The degree of the fitness advantage suggests active inhibition or killing of the ancestor. This may be due to derepression of rhamnolipid production following the inactivation of *mexT* in those populations (Frando et al., 2025; Köhler et al., 2001; Kostylev et al., 2023). Rhamnolipids can inhibit adhesion of competing microbes to surfaces (Boles et al., 2005; Nickzad & Déziel, 2014; Wood et al., 2018) and lyse cells (van Gennip et al., 2009), each of which would increase the competitive advantage in the anoxic biofilm niche (Trejo-Hernández et al., 2014).

Taken together, this study demonstrates that oxygen availability plays a fundamental but understudied role in *P. aeruginosa* resistance evolution. Using experimental evolution under infection-relevant conditions (anoxic growth, biofilm lifestyle, and subinhibitory antibiotic exposure), we identified adaptive pathways that have not been appreciated in standard aerobic laboratory conditions. The convergent selection for mutations in *mexT* across conditions highlights it as a central adaptive locus linking antibiotic resistance, virulence factor derepression, and anoxic niche specialization. The broader patterns of condition-dependent resistance alleles, anoxic-specific biofilm increase, T4P loss reinforcing a sessile lifestyle, and superior anoxic fitness closely parallels the adaptive pathway of *P. aeruginosa* during chronic CF infection (Ciofu & Tolker-Nielsen, 2019; Folkesson et al., 2012). Finally, the anoxic fitness advantages and biofilm phenotypes identified argue for testing and developing therapeutic strategies targeting *P. aeruginosa* anaerobic growth and physiology.

## Methods

### Strains and media

The evolution experiment used *Pseudomonas aeruginosa* MPAO1. Both *P. aeruginosa* MPAO1 *lacZ* positive and MPAO1 *lacZ* negative were utilized to easily identify contamination between evolutionary replicates and aid in phenotypic assays. The minimal media, referred to as M9+, used throughout the experiment contained M9 salts (26 mM Na_2_HPO_4_· 7H_2_O, 22 mM KH_2_PO_4_, 8.6 mM NaCl, 18.7 mM NH_4_Cl, 2.0 mM MgSO_4_, 0.1 mM CaCl_2_), 10 mM glucose, minimum essential medium (MEM) essential amino acids solution (50X,Gibco) (20 mL/liter), MEM nonessential amino acids solution (100X, Gibco) (10 mL/liter), and trace elements A, B, and C (1000X, Corning) (1 mL/liter). 10 mM lactate was also added to the minimal media to approximate some of the nutrient conditions that occur in the cystic fibrosis lung (Palmer et al., 2007). Additionally, 40mM KNO_3_ was added to anaerobic media to support the anaerobic respiration of nitrate. 18mm x 150mm glass tubes were used for all culture conditions with 5 mL of minimal media. Each anaerobic tube was sealed with a butyl rubber stopper (Chemglass), an aluminum seal, and sparged with nitrogen gas to remove oxygen. All cultures were incubated in a roller drum (∼20 rpm) for 24 hours at 37°C.

To aid in confirmation of phenotypic effects of mutations identified, transposon insertion mutants from the *P. aeruginosa* PAO1 Two-Allele Transposon Mutant Library (Held et al., 2012) were obtained through the University of Washington. Strains used included PW1223 [*mexT*-E04::ISlacZ/hah], PW5177 [*mexT*-F04::ISlacZ/hah], PW1728 [*pilT*-E12::ISphoA/hah], PW8623 [*pilB*-G07::ISlacZ/hah], PW6238 [*fimV*-F01::ISphoA/hah], and PW9465 [*pilQ*-B06::ISphoA/hah]. Mutant strains were revived and assayed under the same conditions as the evolved populations described above.

### Evolution experiment

To start the evolution experiment, one colony representing a single clone of *P. aeruginosa* MPAO1 was resuspended in phosphate-buffered saline (PBS) and aliquoted into evolutionary replicate lineages (36 replicates). *Pseudomonas aeruginosa* MPAO1 *lacZ-*positive was used for 18 lineages and *P. aeruginosa* MPAO1 *lacZ-*negative was used for the other 18 lineages, alternating within each condition to control for contamination. For the first half of the experiment (15 days), the antibiotic-exposed replicates were propagated under subinhibitory TOB exposure (0.5xMIC or 0.5 mg/L).

TOB concentration was increased to 1.0 mg/L (1xMIC) for the second half of the experiment (days 16-30). The replicates were also propagated with either biofilm or planktonic selection and oxic or anoxic exposure. The following conditions were tested with six replicates of each: oxic biofilm with TOB, anoxic biofilm with TOB, oxic planktonic with TOB, and anoxic planktonic with TOB. Conditions without TOB exposure were tested with three replicates of each: oxic biofilm, anoxic biofilm, oxic planktonic, anoxic planktonic. Planktonic populations were transferred every day with a 1:100 dilution, adding 50 µL of culture to 5 mL of fresh media. For biofilm populations, a 0.25” PTFE bead (US Plastic Corp.) was transferred to a new tube with fresh media and three new, sterile beads. This allowed for dispersal of bacterial growth from the old bead to colonize the new beads (Poltak & Cooper, 2011). To ensure 24 hours of bacterial growth, beads were marked with either green or white and used on alternating days. An additional planktonic tube, without *P. aeruginosa*, was propagated daily with M9+ media to serve as a negative control. Population samples were saved on days 1, 2, 6, 10, 15, 16, 22, 25, and 30 and frozen at-80°C in 18% glycerol. Planktonic populations were saved by freezing 1mL of liquid culture and biofilm populations were saved following sonication of beads in 1mL PBS.

### Minimum inhibitory concentrations (MICs)

TOB MICs were determined through broth microdilution in both Mueller-Hinton broth and the experimental media (M9+ media) using two-fold increasing concentrations of TOB with a range of 0.125 mg/L to 64 mg/L. MICs were performed according to guidelines set by the Clinical and Laboratory Standards Institute (Clinical and Laboratory Standards Institute, 2026). Freezer stocks of populations were revived in liquid culture into the media and condition in which they were evolved, and following incubation at 37°C, the population was diluted to a 0.5 McFarland standard in minimal media. Biofilm populations were revived into tubes with media and sterile beads. After overnight incubation, a bead was added to 1.0mL of M9+, sonicated and the resuspension was adjusted to a 0.5 McFarland standard if necessary. The dilutions (final concentration of approximately 5×10^5^ CFU/mL) were then inoculated into round bottom 96 well plates with increasing two-fold dilutions of TOB and incubated at 37°C for 16-20 hours. The MIC was then determined as the first concentration at which there was no visible growth. MICs were measured in triplicate for each evolutionary replicate. TOB MICs in M9+ media and Mueller Hinton broth for both *P. aeruginosa* MPAO1 and *Acinetobacter baumannii* ATCC 17978 were used as control on each plate. Anoxic MICs were performed using anaerobic media and placing the assay plate into an anaerobic jar with an AnaeroPack gas generator (Thermo Scientific Mitsubishi).

### Biofilm assays

Biofilm growth was measured for evolved populations using a crystal violet assay (O’Toole, 2011). Freezer stocks of populations were revived in liquid culture into the media and condition in which they were evolved and incubated overnight at 37°C. Cultures were then diluted in fresh minimal media for inoculation into round-bottom 96-well plates. Planktonic populations were diluted 1:100 into media and a bead from the overnight biofilm cultures was added to 1mL of media and sonicated to resuspend the bacterial cells. 100 µL of each dilution was added to a 96-well plate and incubated for 24 hours at 37°C. The following day, the plates were dumped out and rinsed twice with water. 125 µL of 0.1% crystal violet solution was added to each well and the plate was incubated at room temperature for 15 minutes. Following 3-4 rinses with water, the plate was allowed to dry overnight. To solubilize the crystal violet and quantify the biofilm, 125 µL of 30% acetic acid in water was added to each well. The plate was then incubated at room temperature for 15 minutes, and the biofilm was quantified by measuring OD_550_. 30% acetic acid in water was used as a blank and *P. aeruginosa* MPAO1 was used as a control. Six replicates were performed per evolved population.

### Fitness assays

Competitive fitness was measured by co-culturing an evolved population with the oppositely *lac*-marked *P. aeruginosa* MPAO1 ancestor and quantifying changes in frequency. Freezer stocks of populations were revived in liquid culture into the media and condition in which they were evolved and incubated overnight at 37°C. The MPAO1 ancestor was revived from frozen into the experimental M9+ media and incubated overnight at 37°C. The following day, OD_600_ was measured for both the ancestor and evolved population cultures, and they were adjusted to same cell density, diluting with fresh media if necessary. 50 µL of both competitors was then added to a tube with 5 mL fresh media, mixed, a small aliquot was taken for plating to quantify CFU/mL (Time_0_), and the tubes were incubated in roller drum at 37°C overnight. The next day, a transfer was done taking 50 µL of the competition culture, adding it to a new tube with 5 mL of fresh media, and incubating in a roller drum at 37°C overnight. On the final day of the experiment, a small amount of the co-culture was used to measure CFU/mL and frequency of each competitor (Time_48_). CFU/mL were plated on tryptic soy agar (TSA) plates supplemented with 5-bromo-4-cloro-3-indolyl-β-D-galactopyranoside (X-gal) to differentiate between *lacZ-*positive and *lacZ-*negative colonies. Competitions for biofilm populations were performed by alternating green and white beads to ensure the correct bead was being transferred or taken for plating. Beads for quantifying population sizes were added to 1.0 mL PBS, sonicated, and the supernatant was used for quantifying CFU/mL. Anoxic competitions were sparged with nitrogen gas following transfers and prior to overnight incubation.

Frequency of each competitor through time was then determined by counting blue vs. white colonies on Time_0_ and Time_48_ plates and calculating their initial and final population sizes in CFU/mL. Selection rate (r) was calculated using

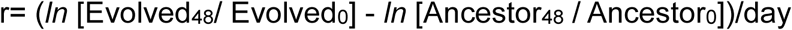

where evolved and ancestor represent the two competitors and the subscripts of 48 and 0 represent the CFU/mL of each competitor at Time_48_ and Time_0_. Once calculated, this ratio represents if either competitor was able to outcompete the other under the conditions within the experiment, with a selection rate of less than zero indicating the ancestor is more fit and a selection rate of greater than zero indicating the evolved population is more fit.

### Twitching motility

To assess type IV pilus-dependent motility across a solid surface, freezer stocks of saved populations were revived in liquid media into the condition in which they were evolved and incubated in a roller drum at 37°C overnight. The following day, the tip of a 10 µL pipette tip was dipped in the overnight culture and stab inoculated through LB agar to the bottom of the petri dish (Darzins, 1993; Déziel et al., 2001). Planktonic-evolved populations were mixed well and inoculated while biofilm-evolved populations required adding a bead to 1.0 mL of PBS, sonicating, and using the resulting resuspension for inoculation. LB agar plates were 1% agar and poured thin (3 mm). Each LB agar petri dish had three replicates of the evolved population being tested as well as an ancestor control (*Pseudomonas aeruginosa* MPAO1) and negative control (*Acinetobacter baumannii* ATCC 17978). The negative control displayed no twitching motility while the ancestor provided a positive control of twitching motility as well as a comparison to the motility of the evolved population. Inoculated plates were then incubated face-up for 48 hours at 37°C. The agar was then carefully lifted out and removed, leaving behind zones of motility and growth on the bottom of the petri dish surrounding each stab. For more effective visualization, the plates were stained with 1% [wt/vol] crystal violet solution, rinsed with tap water, and allowed to dry. To quantify twitching motility, the diameter of the crystal violet stained zones was measured.

### Genome sequencing and analysis

Populations were saved throughout the experiment for whole population sequencing. Sequencing was performed on the ancestor clone as well as four evolutionary replicates of each TOB-exposed condition and two replicates of each condition without TOB exposure from day 30. To prepare for sequencing, freezer stocks of populations were revived into liquid culture into the media and condition from which they were evolved and incubated overnight at 37°C. The overnight cultures were then sampled by taking an aliquot of culture for planktonic populations or transferring two biofilm beads to 1.0 mL of PBS, sonicating, and using the resuspension for biofilm populations. The Qiagen DNeasy blood and tissue kit was used for DNA extraction according to the manufacturer’s protocol. The extracted DNA was then sent to the Microbial Genome Sequencing Center for sequencing library preparation and sequencing on an Illumina NextSeq2000.

Trimmomatic (v0.39) was used to trim the resulting sequences using the following criteria: leading, 20; trailing, 20; sliding window, 4:20; minlen, 70 (Bolger et al., 2014). breseq (v0.35.4) was used to call variants with the default parameters and-p flag to denote population analysis (Deatherage & Barrick, 2014) using *P. aeruginosa* PAO1 (GCF_000006765.1). Within those parameters, mutations were called only if they were at a frequency of at least 5% within the population. The breseq program gdtools was then used to contextualize the GenomeDiff file output from breseq (Deatherage & Barrick, 2014). The SUBTRACT operation was used to remove mutations already present in the ancestor genome from the evolved population mutation output. The ANNOTATE command was then used to annotate the output. Reported mutations were annotated with PAO1 (GCF_000006765.1), but all mutations were cross referenced by following the same mutation-calling procedures with the MPAO1 reference genome (GCF_016107485.1). Reported mutations are available in **Table S1**.

## Data visualization

Data manipulation was done in R (v4.3.1: https://www.r-project.org), using ggplot2 (v3.4.3: https://ggplot2.tidyverse.org) for plotting (Wickham, 2016).

## Data availability

All sequence data can be found in the NCBI SRA under BioProject PRJNA1473505 with accessions SAMN60561751 through SAMN60561810.

## Acknowledgements

This work was funded in part by the Department of the Army, US Army Engineer Research and Development Center award number W9132T-22-2-0001. Strains used in the study were generated by work funded by the Cystic Fibrosis Foundation Grant SINGH24R0.

## References

Abisado, R. G., Kimbrough, J. H., McKee, B. M., Craddock, V. D., Smalley, N. E., Dandekar, A. A., & Chandler, J. R. (2021). Tobramycin Adaptation Enhances Policing of Social Cheaters in Pseudomonas aeruginosa. Applied and Environmental Microbiology, 87(12), e00029–21. 10.1128/AEM.00029-21

Ahmed, M. N., Porse, A., Sommer, M. O. A., Høiby, N., & Ciofu, O. (2018). Evolution of Antibiotic Resistance in Biofilm and Planktonic Pseudomonas aeruginosa Populations Exposed to Subinhibitory Levels of Ciprofloxacin. Antimicrobial Agents and Chemotherapy, 62(8), e00320–18. 10.1128/AAC.00320-18

Andersson, D. I., & Hughes, D. (2010). Antibiotic resistance and its cost: Is it possible to reverse resistance? Nature Reviews. Microbiology, 8(4), 260–271. 10.1038/nrmicro2319

Bjarnsholt, T., Jensen, P. Ø., Fiandaca, M. J., Pedersen, J., Hansen, C. R., Andersen, C. B., Pressler, T., Givskov, M., & Høiby, N. (2009). Pseudomonas aeruginosa biofilms in the respiratory tract of cystic fibrosis patients. Pediatric Pulmonology, 44(6), 547–558. 10.1002/ppul.21011

Bolard, A., Plésiat, P., & Jeannot, K. (2018). Mutations in Gene fusA1 as a Novel Mechanism of Aminoglycoside Resistance in Clinical Strains of Pseudomonas aeruginosa. Antimicrobial Agents and Chemotherapy, 62(2), e01835–17. 10.1128/AAC.01835-17

Boles, B. R., Thoendel, M., & Singh, P. K. (2005). Rhamnolipids mediate detachment of Pseudomonas aeruginosa from biofilms. Molecular Microbiology, 57(5), 1210–1223. 10.1111/j.1365-2958.2005.04743.x

Bolger, A. M., Lohse, M., & Usadel, B. (2014). Trimmomatic: A flexible trimmer for Illumina sequence data. Bioinformatics, 30(15), 2114–2120. 10.1093/bioinformatics/btu170

Borriello, G., Werner, E., Roe, F., Kim, A. M., Ehrlich, G. D., & Stewart, P. S. (2004). Oxygen limitation contributes to antibiotic tolerance of Pseudomonas aeruginosa in biofilms. Antimicrobial Agents and Chemotherapy, 48(7), 2659–2664. 10.1128/AAC.48.7.2659-2664.2004

Burrows, L. L. (2012). Pseudomonas aeruginosa twitching motility: Type IV pili in action. Annual Review of Microbiology, 66, 493–520. 10.1146/annurev-micro-092611-150055

Centers for Disease Control and Prevention (U.S.). (2019). Antibiotic resistance threats in the United States, 2019. Centers for Disease Control and Prevention (U.S.). 10.15620/cdc:82532

Ciofu, O., & Tolker-Nielsen, T. (2019). Tolerance and Resistance of Pseudomonas aeruginosa Biofilms to Antimicrobial Agents-How P. aeruginosa Can Escape Antibiotics. Frontiers in Microbiology, 10, 913. 10.3389/fmicb.2019.00913

Clinical and Laboratory Standards Institute. (2026). M100 | Performance Standards for Antimicrobial Susceptibility Testing. Clinical and Laboratory Standards Institute. https://clsi.org/shop/standards/m100/

Cooper, V. S. (2018). Experimental Evolution as a High-Throughput Screen for Genetic Adaptations. mSphere, 3(3),. 10.1128/msphere.00121-18

Cowley, E. S., Kopf, S. H., LaRiviere, A., Ziebis, W., & Newman, D. K. (2015). Pediatric Cystic Fibrosis Sputum Can Be Chemically Dynamic, Anoxic, and Extremely Reduced Due to Hydrogen Sulfide Formation. mBio, 6(4), e00767–15. 10.1128/mBio.00767-15

Cullen, L., & McClean, S. (2015). Bacterial Adaptation during Chronic Respiratory Infections. Pathogens, 4(1), 66–89. 10.3390/pathogens4010066

Cutruzzolà, F., & Frankenberg-Dinkel, N. (2015). Origin and Impact of Nitric Oxide in Pseudomonas aeruginosa Biofilms. Journal of Bacteriology, 198(1), 55–65. 10.1128/JB.00371-15

Darzins, A. (1993). The pilG gene product, required for Pseudomonas aeruginosa pilus production and twitching motility, is homologous to the enteric, single-domain response regulator CheY. Journal of Bacteriology, 175(18), 5934–5944. 10.1128/jb.175.18.5934-5944.1993

Deatherage, D. E., & Barrick, J. E. (2014). Identification of Mutations in Laboratory-Evolved Microbes from Next-Generation Sequencing Data Using breseq. In L. Sun & W. Shou (Eds.), Engineering and Analyzing Multicellular Systems: Methods and Protocols (pp. 165–188). Springer. 10.1007/978-1-4939-0554-6_12

Déziel, E., Comeau, Y., & Villemur, R. (2001). Initiation of Biofilm Formation byPseudomonas aeruginosa 57RP Correlates with Emergence of Hyperpiliated and Highly Adherent Phenotypic Variants Deficient in Swimming, Swarming, and Twitching Motilities. Journal of Bacteriology, 183(4), 1195–1204. 10.1128/jb.183.4.1195-1204.2001

Figueroa, W., Cazares, A., Ashworth, E. A., Weimann, A., Kadioglu, A., Floto, R. A., & Welch, M. (2025). Mutations in mexT bypass the stringent response dependency of virulence in Pseudomonas aeruginosa. Cell Reports, 44(1). 10.1016/j.celrep.2024.115079

Flemming, H.-C., & Wingender, J. (2010). The biofilm matrix. Nature Reviews. Microbiology, 8(9), 623–633. 10.1038/nrmicro2415

Frando, A., Parsek, R. S., Omar, J., Groleau, M.-C., Trottier, M. C., Smalley, N. E., Déziel, E., & Dandekar, A. A. (2025). Modulation of the Pseudomonas aeruginosa quorum sensing cascade by MexT-regulated factors. mBio, 16(11), e02941–25. 10.1128/mbio.02941-25

Gebhardt, M. J., & Shuman, H. A. (2017). GigA and GigB are Master Regulators of Antibiotic Resistance, Stress Responses, and Virulence in Acinetobacter baumannii. Journal of Bacteriology, 199(10), e00066–17. 10.1128/JB.00066-17

Geller, D. E., Pitlick, W. H., Nardella, P. A., Tracewell, W. G., & Ramsey, B. W. (2002). Pharmacokinetics and bioavailability of aerosolized tobramycin in cystic fibrosis. Chest, 122(1), 219–226. 10.1378/chest.122.1.219

Gloag, E. S., Marshall, C. W., Snyder, D., Lewin, G. R., Harris, J. S., Santos-Lopez, A., Chaney, S. B., Whiteley, M., Cooper, V. S., & Wozniak, D. J. (2019). Pseudomonas aeruginosa Interstrain Dynamics and Selection of Hyperbiofilm Mutants during a Chronic Infection. mBio, 10(4), e01698–19. 10.1128/mBio.01698-19

Global burden of bacterial antimicrobial resistance in 2019: A systematic analysis. (2022). Lancet (London, England), 399(10325), 629–655. 10.1016/S0140-6736(21)02724-0

Gullberg, E., Cao, S., Berg, O. G., Ilbäck, C., Sandegren, L., Hughes, D., & Andersson, D. I. (2011). Selection of Resistant Bacteria at Very Low Antibiotic Concentrations. PLoS Pathogens, 7(7), e1002158. 10.1371/journal.ppat.1002158

Hammond, J. H., Dolben, E. F., Smith, T. J., Bhuju, S., & Hogan, D. A. (2015). Links between Anr and Quorum Sensing in Pseudomonas aeruginosa Biofilms. Journal of Bacteriology, 197(17), 2810–2820. 10.1128/JB.00182-15

Hassett, D. J., Sutton, M. D., Schurr, M. J., Herr, A. B., Caldwell, C. C., & Matu, J. O. (2009). Pseudomonas aeruginosa hypoxic or anaerobic biofilm infections within cystic fibrosis airways. Trends in Microbiology, 17(3), 130–138. 10.1016/j.tim.2008.12.003

Held, K., Ramage, E., Jacobs, M., Gallagher, L., & Manoil, C. (2012). Sequence-Verified Two-Allele Transposon Mutant Library for Pseudomonas aeruginosa PAO1. Journal of Bacteriology, 194(23), 6387–6389. 10.1128/JB.01479-12

Hendricks, M. R., Lashua, L. P., Fischer, D. K., Flitter, B. A., Eichinger, K. M., Durbin, J. E., Sarkar, S. N., Coyne, C. B., Empey, K. M., & Bomberger, J. M. (2016). Respiratory syncytial virus infection enhances Pseudomonas aeruginosa biofilm growth through dysregulation of nutritional immunity. Proceedings of the National Academy of Sciences of the United States of America, 113(6), 1642–1647. 10.1073/pnas.1516979113

Hill, D., Rose, B., Pajkos, A., Robinson, M., Bye, P., Bell, S., Elkins, M., Thompson, B., MacLeod, C., Aaron, S. D., & Harbour, C. (2005). Antibiotic Susceptibilities of Pseudomonas aeruginosa Isolates Derived from Patients with Cystic Fibrosis under Aerobic, Anaerobic, and Biofilm Conditions. Journal of Clinical Microbiology, 43(10), 5085–5090. 10.1128/JCM.43.10.5085-5090.2005

Høiby, N., Bjarnsholt, T., Givskov, M., Molin, S., & Ciofu, O. (2010). Antibiotic resistance of bacterial biofilms. International Journal of Antimicrobial Agents, 35(4), 322–332. 10.1016/j.ijantimicag.2009.12.011

Köhler, T., Michéa-Hamzehpour, M., Henze, U., Gotoh, N., Curty, L. K., & Pechère, J. C. (1997). Characterization of MexE-MexF-OprN, a positively regulated multidrug efflux system of Pseudomonas aeruginosa. Molecular Microbiology, 23(2), 345–354. 10.1046/j.1365-2958.1997.2281594.x

Köhler, T., van Delden, C., Curty, L. K., Hamzehpour, M. M., & Pechere, J.-C. (2001). Overexpression of the MexEF-OprN Multidrug Efflux System Affects Cell-to-Cell Signaling in Pseudomonas aeruginosa. Journal of Bacteriology, 183(18), 5213–5222. 10.1128/JB.183.18.5213-5222.2001

Kostylev, M., Smalley, N. E., Chao, M. H., & Greenberg, E. P. (2023). Relationship of the transcription factor MexT to quorum sensing and virulence in Pseudomonas aeruginosa. Journal of Bacteriology, 205(12), e00226–23. 10.1128/jb.00226-23

La Rosa, R., Johansen, H. K., & Molin, S. (2019). Adapting to the Airways: Metabolic Requirements of Pseudomonas aeruginosa during the Infection of Cystic Fibrosis Patients. Metabolites, 9(10), 234. 10.3390/metabo9100234

LaCroix, R. A., Palsson, B. O., & Feist, A. M. (2017). A Model for Designing Adaptive Laboratory Evolution Experiments. Applied and Environmental Microbiology, 83(8), e03115–16. 10.1128/AEM.03115-16

Lau, C. H.-F., Fraud, S., Jones, M., Peterson, S. N., & Poole, K. (2013). Mutational Activation of the AmgRS Two-Component System in Aminoglycoside-Resistant Pseudomonas aeruginosa. Antimicrobial Agents and Chemotherapy, 57(5), 2243–2251. 10.1128/AAC.00170-13

Lee, B., Haagensen, J. A. J., Ciofu, O., Andersen, J. B., Høiby, N., & Molin, S. (2005). Heterogeneity of Biofilms Formed by Nonmucoid Pseudomonas aeruginosa Isolates from Patients with Cystic Fibrosis. Journal of Clinical Microbiology, 43(10), 5247–5255. 10.1128/JCM.43.10.5247-5255.2005

Lee, S., Gallagher, L., & Manoil, C. (2021). Reconstructing a Wild-Type Pseudomonas aeruginosa Reference Strain PAO1. Journal of Bacteriology, 203(14), e0017921. 10.1128/JB.00179-21

Lee, S., Hinz, A., Bauerle, E., Angermeyer, A., Juhaszova, K., Kaneko, Y., Singh, P. K., & Manoil, C. (2009). Targeting a bacterial stress response to enhance antibiotic action. Proceedings of the National Academy of Sciences of the United States of America, 106(34), 14570–14575. 10.1073/pnas.0903619106

Leighton, T. L., Buensuceso, R. N. C., Howell, P. L., & Burrows, L. L. (2015). Biogenesis of Pseudomonas aeruginosa type IV pili and regulation of their function. Environmental Microbiology, 17(11), 4148–4163. 10.1111/1462-2920.12849

Lieberman, T. D., Michel, J.-B., Aingaran, M., Potter-Bynoe, G., Roux, D., Davis, M. R., Skurnik, D., Leiby, N., LiPuma, J. J., Goldberg, J. B., McAdam, A. J., Priebe, G. P., & Kishony, R. (2011). Parallel bacterial evolution within multiple patients identifies candidate pathogenicity genes. Nature Genetics, 43(12), 1275–1280. 10.1038/ng.997

Llanes, C., Köhler, T., Patry, I., Dehecq, B., van Delden, C., & Plésiat, P. (2011). Role of the MexEF-OprN Efflux System in Low-Level Resistance of Pseudomonas aeruginosa to Ciprofloxacin ▿. Antimicrobial Agents and Chemotherapy, 55(12), 5676–5684. 10.1128/AAC.00101-11

López-Causapé, C., Sommer, L. M., Cabot, G., Rubio, R., Ocampo-Sosa, A. A., Johansen, H. K., Figuerola, J., Cantón, R., Kidd, T. J., Molin, S., & Oliver, A. (2017). Evolution of the Pseudomonas aeruginosa mutational resistome in an international Cystic Fibrosis clone. Scientific Reports, 7, 5555. 10.1038/s41598-017-05621-5

Luong, P. M., Shogan, B. D., Zaborin, A., Belogortseva, N., Shrout, J. D., Zaborina, O., & Alverdy, J. C. (2014). Emergence of the P2 Phenotype in Pseudomonas aeruginosa PAO1 Strains Involves Various Mutations in mexT or mexF. Journal of Bacteriology, 196(2), 504–513. 10.1128/JB.01050-13

Lyczak, J. B., Cannon, C. L., & Pier, G. B. (2002). Lung infections associated with cystic fibrosis. Clinical Microbiology Reviews, 15(2), 194–222. 10.1128/CMR.15.2.194-222.2002

Martin, I., Waters, V., & Grasemann, H. (2021). Approaches to Targeting Bacterial Biofilms in Cystic Fibrosis Airways. International Journal of Molecular Sciences, 22(4), 2155. 10.3390/ijms22042155

Martin, L. W., Gray, A. R., Brockway, B., & Lamont, I. L. (2023). Pseudomonas aeruginosa is oxygen-deprived during infection in cystic fibrosis lungs, reducing the effectiveness of antibiotics. FEMS Microbiology Letters, 370, fnad076. 10.1093/femsle/fnad076

Marvig, R. L., Sommer, L. M., Molin, S., & Johansen, H. K. (2015). Convergent evolution and adaptation of Pseudomonas aeruginosa within patients with cystic fibrosis. Nature Genetics, 47(1), 57–64. 10.1038/ng.3148

Masuda, N., Sakagawa, E., Ohya, S., Gotoh, N., Tsujimoto, H., & Nishino, T. (2000). Substrate Specificities of MexAB-OprM, MexCD-OprJ, and MexXY-OprM Efflux Pumps in Pseudomonas aeruginosa. Antimicrobial Agents and Chemotherapy, 44(12), 3322–3327. 10.1128/aac.44.12.3322-3327.2000

Melnyk, A. H., Wong, A., & Kassen, R. (2015). The fitness costs of antibiotic resistance mutations. Evolutionary Applications, 8(3), 273–283. 10.1111/eva.12196

Mogayzel, P. J., Naureckas, E. T., Robinson, K. A., Mueller, G., Hadjiliadis, D., Hoag, J. B., Lubsch, L., Hazle, L., Sabadosa, K., Marshall, B., & Pulmonary Clinical Practice Guidelines Committee. (2013). Cystic fibrosis pulmonary guidelines. Chronic medications for maintenance of lung health. American Journal of Respiratory and Critical Care Medicine, 187(7), 680–689. 10.1164/rccm.201207-1160oe

Moser, C., Jensen, P. Ø., Thomsen, K., Kolpen, M., Rybtke, M., Lauland, A. S., Trøstrup, H., & Tolker-Nielsen, T. (2021). Immune Responses to Pseudomonas aeruginosa Biofilm Infections. Frontiers in Immunology, 12, 625597. 10.3389/fimmu.2021.625597

Naghavi, M., Vollset, S. E., Ikuta, K. S., Swetschinski, L. R., Gray, A. P., Wool, E. E., Aguilar, G. R., Mestrovic, T., Smith, G., Han, C., Hsu, R. L., Chalek, J., Araki, D. T., Chung, E., Raggi, C., Hayoon, A. G., Weaver, N. D., Lindstedt, P. A., Smith, A. E.,…Murray, C. J. L. (2024). Global burden of bacterial antimicrobial resistance 1990–2021: A systematic analysis with forecasts to 2050. The Lancet, 404(10459), 1199–1226. 10.1016/S0140-6736(24)01867-1

Nickzad, A., & Déziel, E. (2014). The involvement of rhamnolipids in microbial cell adhesion and biofilm development—An approach for control? Letters in Applied Microbiology, 58(5), 447–453. 10.1111/lam.12211

O’Toole, G. A. (2011). Microtiter Dish Biofilm Formation Assay. Journal of Visualized Experiments (JoVE), (47), e2437. 10.3791/2437

Palmer, K. L., Aye, L. M., & Whiteley, M. (2007). Nutritional Cues Control Pseudomonas aeruginosa Multicellular Behavior in Cystic Fibrosis Sputum. Journal of Bacteriology, 189(22), 8079–8087. 10.1128/jb.01138-07

Park, J.-S., Choi, H.-Y., & Kim, W.-G. (2020). The Nitrite Transporter Facilitates Biofilm Formation via Suppression of Nitrite Reductase and Is a New Antibiofilm Target in Pseudomonas aeruginosa. mBio, 11(4), e00878–20. 10.1128/mBio.00878-20

Pereira, C., Warsi, O. M., & Andersson, D. I. (2023). Pervasive Selection for Clinically Relevant Resistance and Media Adaptive Mutations at Very Low Antibiotic Concentrations. Molecular Biology and Evolution, 40(1), msad010. 10.1093/molbev/msad010

Pflüger-Grau, K., & Görke, B. (2010). Regulatory roles of the bacterial nitrogen-related phosphotransferase system. Trends in Microbiology, 18(5), 205–214. 10.1016/j.tim.2010.02.003

Poltak, S. R., & Cooper, V. S. (2011). Ecological succession in long-term experimentally evolved biofilms produces synergistic communities. The ISME Journal, 5(3), 369–378. 10.1038/ismej.2010.136

Ramsay, K. A., McTavish, S. M., Wardell, S. J. T., & Lamont, I. L. (2021). The Effects of Sub-inhibitory Antibiotic Concentrations on Pseudomonas aeruginosa: Reduced Susceptibility Due to Mutations. Frontiers in Microbiology, 12, 789550. 10.3389/fmicb.2021.789550

Ramsey, B. W., Pepe, M. S., Quan, J. M., Otto, K. L., Montgomery, A. B., Williams-Warren, J., Vasiljev-K, M., Borowitz, D., Bowman, C. M., Marshall, B. C., Marshall, S., & Smith, A. L. (1999). Intermittent administration of inhaled tobramycin in patients with cystic fibrosis. Cystic Fibrosis Inhaled Tobramycin Study Group. The New England Journal of Medicine, 340(1), 23–30. 10.1056/NEJM199901073400104

Richardot, C., Juarez, P., Jeannot, K., Patry, I., Plésiat, P., & Llanes, C. (2016). Amino Acid Substitutions Account for Most MexS Alterations in Clinical nfxC Mutants of Pseudomonas aeruginosa. Antimicrobial Agents and Chemotherapy, 60(4), 2302–2310. 10.1128/AAC.02622-15

Rodnina, M. V. (2013). Translocation in Action. Science, 340(6140), 1534–1535. 10.1126/science.1240090

Römling, U., Galperin, M. Y., & Gomelsky, M. (2013). Cyclic di-GMP: The First 25 Years of a Universal Bacterial Second Messenger. Microbiology and Molecular Biology Reviews: MMBR, 77(1), 1–52. 10.1128/MMBR.00043-12

Rosenthal, V. D., Al-Abdely, H. M., El-Kholy, A. A., AlKhawaja, S. A. A., Leblebicioglu, H., Mehta, Y., Rai, V., Hung, N. V., Kanj, S. S., Salama, M. F., Salgado-Yepez, E., Elahi, N., Morfin Otero, R., Apisarnthanarak, A., De Carvalho, B. M., Ider, B. E., Fisher, D., Buenaflor, M. C. S. G., Petrov, M. M.,…Remaining authors. (2016). International Nosocomial Infection Control Consortium report, data summary of 50 countries for 2010-2015: Device-associated module. American Journal of Infection Control, 44(12), 1495–1504. 10.1016/j.ajic.2016.08.007

Rouillard, K. R., Esther, C. P., Kissner, W. J., Plott, L. M., Bowman, D. W., Markovetz, M. R., & Hill, D. B. (2024). Combination treatment to improve mucociliary transport of Pseudomonas aeruginosa biofilms. PLOS ONE, 19(2), e0294120. 10.1371/journal.pone.0294120

Santos-Lopez, A., Marshall, C. W., Scribner, M. R., Snyder, D. J., & Cooper, V. S. (2019). Evolutionary pathways to antibiotic resistance are dependent upon environmental structure and bacterial lifestyle. eLife, 8, e47612. 10.7554/eLife.47612

Schurek, K. N., Marr, A. K., Taylor, P. K., Wiegand, I., Semenec, L., Khaira, B. K., & Hancock, R. E. W. (2008). Novel genetic determinants of low-level aminoglycoside resistance in Pseudomonas aeruginosa. Antimicrobial Agents and Chemotherapy, 52(12), 4213–4219. 10.1128/AAC.00507-08

Scribner, M. R., Santos-Lopez, A., Marshall, C. W., Deitrick, C., & Cooper, V. S. (2020). Parallel Evolution of Tobramycin Resistance across Species and Environments. mBio, 11(3), e00932–20. 10.1128/mBio.00932-20

Secor, P. R., Jennings, L. K., Michaels, L. A., Sweere, J. M., Singh, P. K., Parks, W. C., & Bollyky, P. L. (2015). Biofilm assembly becomes crystal clear – filamentous bacteriophage organize the Pseudomonas aeruginosa biofilm matrix into a liquid crystal. Microbial Cell, 3(1), 49–52. 10.15698/mic2016.01.475

Sherrard, L. J., Tai, A. S., Wee, B. A., Ramsay, K. A., Kidd, T. J., Ben Zakour, N. L., Whiley, D. M., Beatson, S. A., & Bell, S. C. (2017). Within-host whole genome analysis of an antibiotic resistant Pseudomonas aeruginosa strain sub-type in cystic fibrosis. PloS One, 12(3), e0172179. 10.1371/journal.pone.0172179

Smith, E. E., Buckley, D. G., Wu, Z., Saenphimmachak, C., Hoffman, L. R., D’Argenio, D. A., Miller, S. I., Ramsey, B. W., Speert, D. P., Moskowitz, S. M., Burns, J. L., Kaul, R., & Olson, M. V. (2006). Genetic adaptation by Pseudomonas aeruginosa to the airways of cystic fibrosis patients. Proceedings of the National Academy of Sciences of the United States of America, 103(22), 8487–8492. 10.1073/pnas.0602138103

Soares, A., Alexandre, K., & Etienne, M. (2020). Tolerance and Persistence of Pseudomonas aeruginosa in Biofilms Exposed to Antibiotics: Molecular Mechanisms, Antibiotic Strategies and Therapeutic Perspectives. Frontiers in Microbiology, 11, 2057. 10.3389/fmicb.2020.02057

Sommer, M. O. A., Munck, C., Toft-Kehler, R. V., & Andersson, D. I. (2017). Prediction of antibiotic resistance: Time for a new preclinical paradigm? Nature Reviews Microbiology, 15(11), 689–696. 10.1038/nrmicro.2017.75

Stewart, P. S., & Costerton, J. W. (2001). Antibiotic resistance of bacteria in biofilms. Lancet, 358(9276), 135–138. 10.1016/s0140-6736(01)05321-1

Tian, Z.-X., Fargier, E., Mac Aogáin, M., Adams, C., Wang, Y.-P., & O’Gara, F. (2009). Transcriptome profiling defines a novel regulon modulated by the LysR-type transcriptional regulator MexT in Pseudomonas aeruginosa. Nucleic Acids Research, 37(22), 7546–7559. 10.1093/nar/gkp828

Tian, Z.-X., Mac Aogáin, M., O’Connor, H. F., Fargier, E., Mooij, M. J., Adams, C., Wang, Y.-P., & O’Gara, F. (2009). MexT modulates virulence determinants in *Pseudomonas aeruginosa* independent of the MexEF-OprN efflux pump. Microbial Pathogenesis, 47(4), 237–241. 10.1016/j.micpath.2009.08.003

Toyofuku, M., Uchiyama, H., & Nomura, N. (2012). Social Behaviours under Anaerobic Conditions in Pseudomonas aeruginosa. International Journal of Microbiology, 2012, 405191. 10.1155/2012/405191

Traverse, C. C., Mayo-Smith, L. M., Poltak, S. R., & Cooper, V. S. (2013). Tangled bank of experimentally evolved Burkholderia biofilms reflects selection during chronic infections. Proceedings of the National Academy of Sciences of the United States of America, 110(3), E250–259. 10.1073/pnas.1207025110

Trejo-Hernández, A., Andrade-Domínguez, A., Hernández, M., & Encarnación, S. (2014). Interspecies competition triggers virulence and mutability in Candida albicans–Pseudomonas aeruginosa mixed biofilms. The ISME Journal, 8(10), 1974–1988. 10.1038/ismej.2014.53

van der Horst, M. A., Schuurmans, J. M., Smid, M. C., Koenders, B. B., & ter Kuile, B. H. (2011). De novo acquisition of resistance to three antibiotics by Escherichia coli. Microbial Drug Resistance, 17(2), 141–147. 10.1089/mdr.2010.0101

van Gennip, M., Christensen, L. D., Alhede, M., Phipps, R., Jensen, P. Ø., Christophersen, L., Pamp, S. J., Moser, C., Mikkelsen, P. J., Koh, A. Y., Tolker-Nielsen, T., Pier, G. B., Høiby, N., Givskov, M., & Bjarnsholt, T. (2009). Inactivation of the rhlA gene in Pseudomonas aeruginosa prevents rhamnolipid production, disabling the protection against polymorphonuclear leukocytes. *APMIS: Acta Pathologica*, Microbiologica, et Immunologica Scandinavica, 117(7), 537–546. 10.1111/j.1600-0463.2009.02466.x

Walters, M. C., Roe, F., Bugnicourt, A., Franklin, M. J., & Stewart, P. S. (2003). Contributions of antibiotic penetration, oxygen limitation, and low metabolic activity to tolerance of Pseudomonas aeruginosa biofilms to ciprofloxacin and tobramycin. Antimicrobial Agents and Chemotherapy, 47(1), 317–323. 10.1128/AAC.47.1.317-323.2003

Wehbi, H., Portillo, E., Harvey, H., Shimkoff, A. E., Scheurwater, E. M., Howell, P. L., & Burrows, L. L. (2011). The Peptidoglycan-Binding Protein FimV Promotes Assembly of the Pseudomonas aeruginosa Type IV Pilus Secretin. Journal of Bacteriology, 193(2), 540– 550. 10.1128/JB.01048-10

WHO Bacterial Priority Pathogens List 2024: Bacterial Pathogens of Public Health Importance, to Guide Research, Development, and Strategies to Prevent and Control Antimicrobial Resistance (1st ed). (2024). World Health Organization.

Wickham, H. (2016). ggplot2: Elegant Graphics for Data Analysis. Springer-Verlag.

Wistrand-Yuen, E., Knopp, M., Hjort, K., Koskiniemi, S., Berg, O. G., & Andersson, D. I. (2018). Evolution of high-level resistance during low-level antibiotic exposure. Nature Communications, 9, 1599. 10.1038/s41467-018-04059-1

Wood, T. L., Gong, T., Zhu, L., Miller, J., Miller, D. S., Yin, B., & Wood, T. K. (2018). Rhamnolipids from Pseudomonas aeruginosa disperse the biofilms of sulfate-reducing bacteria. NPJ Biofilms and Microbiomes, 4, 22. 10.1038/s41522-018-0066-1

Worlitzsch, D., Tarran, R., Ulrich, M., Schwab, U., Cekici, A., Meyer, K. C., Birrer, P., Bellon, G., Berger, J., Weiss, T., Botzenhart, K., Yankaskas, J. R., Randell, S., Boucher, R. C., & Döring, G. (2002). Effects of reduced mucus oxygen concentration in airway Pseudomonas infections of cystic fibrosis patients. The Journal of Clinical Investigation, 109(3), 317– 325. 10.1172/JCI13870

